# Alterations in the Antioxidant Metabolites in Cotton Leaf Curl Disease Infected and Healthy Cotton Leaves

**DOI:** 10.1101/2021.11.10.468097

**Authors:** Atiq Ur Rehman, Muhammad Anjum Zia, Muhammad Mubin, Muhammad Shahid

## Abstract

The upland cotton (*Gossypium hirsutum*) belongs to the family *Malvaceae* and cultivated since ancient times in warmer parts of the world. Pakistan is the 4^th^ largest cotton producing country and prominent cotton yarn exporter of the world. Cotton is the major source of natural fiber and significantly contributes to the economy of Pakistan. There are many factors (biotic and abiotic) affecting the cotton yield in Pakistan. The cotton leaf curl disease (CLCuD) is one of the biotic factors and endemic in the most cotton growing areas of the country. The antioxidant enzymes and antioxidant biochemical metabolites play important role during stress. The present study was planned to compare the antioxidant enzymes and metabolites from healthy and CLCuD infected cotton leaves. Antioxidant enzymes activities including catalase, peroxidase and peroxidase were measured through different antioxidant assays and different antioxidant metabolites were also determined. During stress condition antioxidant metabolites serve as signal for the activation of antioxidant enzymes. Present study revealed that antioxidant enzymes were significantly higher in the infected cotton leaves as compared to the healthy cotton leaves. While the amount of antioxidant metabolites like total phenolic contents (TPC) and total flavonoid contents (TFC) were higher in healthy cotton leaves as compared to the infected leaves.

## Introduction

The upland cotton belongs to the family Malvaceae and cultivated since ancient times in warmer parts of the world. The crop is mainly cultivated for its precious fiber but it also contributes 80% share to oil production of Pakistan in addition to its lint [1]. Pakistan is the 4^th^ largest cotton producing country of the world and prominent cotton yarn exporter of the world. The cotton crop is considered backbone of national economy as it is cultivated on the area of 3 million hectares in the country [2]. The economy of Pakistan greatly dependents on cotton and cotton products. It has less than 10% share in agriculture value but provide about 2% share to the GDP of the country [3]. It is economically most important crop of the country contributing up to 60 % of total exports and having largest value chain employing 50% of total industrial labor from field to the final valuable product [4]. The single crop provides the livelihood to the millions of farmers, agricultural labor, industrial labor and traders [5]. There are multiple biotic and abiotic factors that affect the yield of cotton. Among biotic factors, insect pests and plant diseases are leading challenges to the sustainable cotton cultivation. In Indian subcontinent especially the Pakistan “the cotton leaf curl disease” is the major dilemma to cotton crop since last few decades. The disease also deteriorate the quality of fiber by affecting traits like percentage of ginning out turn, fiber fineness, staple length, fiber uniformity index, fiber bundle strength and maturity ratio due to the changes in metabolites (cellulose, pectin, waxes and proteins) composition in fiber in addition to the effect on total crop yield [6].

The reactive oxygen species (ROS) are not always harmful and toxic for the plants but also perform special assigned role during development and regulation of metabolism of plant. ROS may also act as indispensable signal molecules in plant biological system such as in response to high light [7], cell cycle regulation [8], germination of seed [9], development of root system [10] and during pathogenic infection [11]. The aerobic organisms to counteract oxidative stress caused by ROS produce antioxidant enzymes and antioxidant compounds. ROS comprises of molecular oxygen (^·^O^−^), superoxide radical (^·^O2^−^), hydroxyl radical (^·^OH) and hydrogen peroxide (H_2_O_2_) produced during oxidation/reduction (redox) reactions of aerobic metabolism as bi-products [12]. To detoxify ROS, there are three antioxidant enzymes with key roles in plants. The superoxide dismutase (SOD) converts ^·^O_2_^−^ to O_2_ and H_2_O_2_. The H_2_O_2_ is then degraded by catalase and peroxidases [13]. Catalases have higher rate of reaction and low affinity to H_2_O_2_ as compared to that of peroxidases which have higher affinity and can even detoxify H_2_O_2_ in lower concentrations [14].

Phenolic metabolites are important plant constituents including phenolic acids, flavonoids, stilbenes, lignins, lignans and tannins etc. [15]. These compounds are very good electron donors and contribute their role as antioxidants [16]. Flavonoids are versatile secondary metabolites with polyphenolic molecular structure and low molecular weight. They are widely distributed in plants and perform diverse physiological functions in plants especially under biotic and abiotic stress. The diverse functions perform by these biomolecules are protective role against salt stress [17], drought stress [18], partial detoxification of ROS [19, 20]. Enormous flavonoids have also been characterized with diverse biological roles for example, shield against ultra violet radiation, signaling during plant microbe interaction for nodulation [21], defense against pest and phytopathogens [22], male fertility of plant, production of visual signals by coloring the flowers to attract pollinators [23] and in transportation of hormones from one part of plant to the other. Few additional functions are protection against photo-radiation and enhancement of nutritional retrieval efficiency during senescence [24].

The plant response to elicitors is the developing field of research in plant physiology. Among commonly used elicitors are salicylic acid, methyl salicylate, benzothiadiazide, benzoic acid and chitosan etc. which activate different defense related enzymes and alter the production of phenolic compounds [25]. These compounds stimulate the interaction between pathogens and plants that subsequently triggers plant’s defense system which constrain the invasion of pathogenic viruses. The tannins and phenols are well studied examples and these biomolecules control resistance genes regulation [26]. The role of these biochemical metabolites is crucial in infection of CLCuD and thus survival of plant under stress. The study of these important biomolecules can provide useful information regarding host defense mechanism against viral infection. In the present study, we have compared the activity of antioxidant enzymes as well as flavonoids and total phenolic contents in the leaves of healthy cotton plants and CLCuD infected cotton plants.

## Materials and Methods

This research work was carried out at Medicinal Biochemistry Lab, Department of Biochemistry, University of Agriculture, Faisalabad.

### Sample collection

The leaf samples were collected randomly from eight locations of cotton growing areas of Punjab, Pakistan, during field survey in September 2019. Total six sample (three infected and three healthy) from each location were collected. The samples were collected in labeled polyether zipper bags and preserved in thermophore box containing ice bricks. The samples were stored in -70°C freezer for long term storage.

### Quantification of Antioxidant Enzymes

The leaf samples (10 mg) of each plant were taken in 2 mL tubes and 1 ml of pre-cooled 50 MM potassium phosphate buffer was add to the tubes containing leaf samples and samples were grounded using bead beater (Omni bead rupture, Canada). The samples were centrifuged for 10 minutes at 13000 rpm to pellet the leaf contents. The supernatant containing the cell lysate were collected in fresh prelabelled eppendorf tubes.

### Catalase Estimation

The catalase is present in almost every aerobic cell and detoxify the H_2_O_2._ The enzyme activity was determined in the leaf samples using Amplex Red Catalase Assay kit (A22180).

### Reaction Mixture and Standard curve

The reaction mixture was prepared using solution provided in the kit and according to instructions of manufacturer. The 20 MM H_2_O_2_ was prepared by diluting 23 µl of 3 % H_2_O_2_ in 977 µl of deionized water (dH_2_O) and for final use 40 µM H_2_O_2_ was prepared using 10 µl of 20 MM in 490 µl of 1X reaction buffer. The 260 µg Amplex Red reagent was dissolved in 100 µl of dimethylsulfoxide (DMSO) to prepare the working solution of Amplex Red. 100 µl of 1 X reaction buffer was added to the horseradish peroxide (HRP) vail to get 100 U/ml solution. The working reaction mixture of Amplex Red and HRP 100 µM and 400 mU, respectively was prepared from previously prepared solutions. To prepare standards, 100 µl of dH_2_O was added to the vail containing catalase and eight standards ranging from 0 to 4 U/ml were prepared using 1 X reaction buffer. The standard curve was plotted using these standard solutions of catalase (Supplementary Figure S1). 25 µl of each standard was pipetted in three wells of ELISA plate and 25 µl of 40 µM H_2_O_2_ was added to each well using multichannel pipette. The mixture was allowed to react for 3 minutes at 37°C followed by addition of 50 µl Amplex Red reagent and horseradish peroxidase working solution to make the reaction mixture to 100 µl. The plate was incubated at 25 °C for 5 minutes to complete the reaction and the incubation time was optimized after multiple trails.

The absorbance of the reaction plate was measured at 560 nm by using micro well plate reader (BioTek, µ-QuantTM, USA). The plant samples were arranged in the racks according to descending order and kept at room temperature before use. The assay was carried out using 25 µl sample and 25 µl of 40 µM H_2_O_2_ and plate was placed at 37 °C for 3 minutes. Apmlex Red and HRP solution were added and plate was placed at 25 °C for 5 minutes before measurement of absorbance.

### Calculation of Enzyme activity

To calculate the activity of catalase in our samples linear regression equation (Y=-0.0002X + 0.8033) obtained from standard curve was used (Supplementary Figure S1). The U/mL units were converted to the U/g of sample weight by multiplying with dilution factors and expressed in (Table 1). 1 U of catalase is described as the quantity of enzyme which decomposes 1.0 µM H_2_O_2_ per minutes at optimum conditions of pH and temperature.

**Table 1.** Location of collected sample, physical condition of the plant, antioxidant enzymes superoxide dismutase, catalase and peroxidase in unit/g of fresh weight and antioxidant metabolites, total phenolic contents in equivalence of gallic acid (GAE) and total flavonoid contents in equivalence of quercetin (QE) from eight different locations in Punjab, Pakistan.

| Sample No. | Location | Physical condition of cotton sample | Superoxide Dismutase (SOD) (Unit/g of FW) | Catalase (CAT) (Unit/g of FW) | Peroxidases (PODs) (Unit/g of FW) | Total phenolic contents (mg GAE/g. FW) | Total flavonoids contents (mg QE/g. FW) |
| --- | --- | --- | --- | --- | --- | --- | --- |
| 1 | District Faisalabad<br>31°23'53.3"N<br>73°02'01.0"E | healthy | 123.62 | 129.58 | 132.81 | 35.60 | 25.62 |
| 2 |  | healthy | 134.62 | 141.58 | 133.52 | 38.68 | 27.22 |
| 3 |  | healthy | 125.16 | 133.12 | 143.46 | 37.28 | 27.52 |
| 4 |  | infected | 215.76 | 226.53 | 209.49 | 32.10 | 18.42 |
| 5 |  | infected | 216.57 | 228.34 | 210.51 | 35.25 | 16.92 |
| 6 |  | infected | 197.53 | 210.30 | 201.81 | 35.04 | 19.52 |
| 7 | District Multan | healthy | 114.16 | 124.65 | 120.75 | 58.91 | 26.52 |
| 8 | 30°14'55.3"N<br>71°35'20.3"E | healthy | 124.46 | 128.40 | 132.50 | 57.72 | 29.22 |
| 9 |  | healthy | 135.13 | 140.07 | 113.86 | 59.82 | 27.12 |
| 10 |  | infected | 205.37 | 211.31 | 237.45 | 54.08 | 23.32 |
| 11 |  | infected | 217.25 | 224.19 | 244.58 | 58.00 | 21.92 |
| 12 |  | infected | 211.65 | 219.59 | 252.33 | 57.58 | 23.62 |
| 13 | District Multan<br>29°34'33.2"N<br>71°38'51.3"E | healthy | 124.46 | 132.69 | 59.90 | 60.66 | 25.52 |
| 14 |  | healthy | 134.86 | 144.09 | 52.38 | 65.42 | 24.43 |
| 15 |  | healthy | 140.71 | 150.94 | 60.91 | 62.13 | 25.62 |
| 16 |  | infected | 240.47 | 244.63 | 167.20 | 48.13 | 19.98 |
| 17 |  | infected | 245.35 | 250.51 | 164.85 | 60.73 | 20.62 |
| 18 |  | infected | 248.75 | 254.91 | 181.84 | 61.08 | 19.42 |
| 19 | Matila chowk<br>29°59'52.4"N<br>71°49'36.4"E | healthy | 142.94 | 150.10 | 237.45 | 47.15 | 25.82 |
| 20 |  | healthy | 133.46 | 141.62 | 239.41 | 49.67 | 27.42 |
| 21 |  | healthy | 123.80 | 132.96 | 253.97 | 49.81 | 28.22 |
| 22 |  | infected | 205.09 | 215.25 | 264.00 | 38.68 | 23.32 |
| 23 |  | infected | 195.54 | 206.70 | 253.19 | 27.06 | 20.62 |
| 24 |  | infected | 215.31 | 227.47 | 268.78 | 34.20 | 21.92 |
| 25 | District Multan<br>30°06'19.0"N<br>71°37'41.3"E | healthy | 145.36 | 155.96 | 115.97 | 83.62 | 29.22 |
| 26 |  | healthy | 136.54 | 148.14 | 102.74 | 88.66 | 26.82 |
| 27 |  | healthy | 124.26 | 136.86 | 101.17 | 74.66 | 28.42 |
| 28 |  | infected | 228.61 | 240.05 | 248.81 | 26.64 | 14.62 |
| 29 |  | infected | 239.74 | 251.18 | 244.97 | 40.85 | 16.12 |
| 30 |  | infected | 216.86 | 228.30 | 229.23 | 29.79 | 17.32 |
| 31 | District Lodhran<br>29°40'25.2"N<br>71°35'25.1"E | healthy | 123.23 | 133.07 | 241.29 | 38.12 | 27.12 |
| 32 |  | healthy | 142.91 | 151.36 | 222.57 | 34.76 | 29.82 |
| 33 |  | healthy | 133.17 | 142.62 | 225.78 | 48.55 | 27.92 |
| 34 |  | infected | 215.36 | 225.81 | 293.06 | 31.82 | 9.42 |
| 35 |  | infected | 225.13 | 236.58 | 285.22 | 30.91 | 12.02 |
| 36 |  | infected | 232.19 | 241.16 | 300.89 | 21.11 | 8.82 |
| 37 | District Lodhran | healthy | 124.41 | 134.38 | 126.39 | 72.14 | 26.42 |
| 38 | 29°34'53.7"N | healthy | 142.29 | 153.26 | 129.68 | 61.92 | 26.02 |
| 39 | 71°38'09.2"E | healthy | 134.53 | 146.50 | 129.44 | 54.92 | 27.22 |
| 40 |  | infected | 212.48 | 224.78 | 288.44 | 38.82 | 12.22 |
| 41 |  | infected | 236.69 | 243.07 | 286.79 | 32.59 | 10.92 |
| 42 |  | infected | 228.39 | 235.77 | 294.62 | 32.52 | 12.72 |
| 43 | District Jhang<br>30°37'00.9"N<br>71°40'28.5"E | healthy | 124.47 | 131.85 | 107.91 | 68.64 | 52.02 |
| 44 |  | healthy | 135.62 | 144.00 | 116.05 | 73.89 | 51.02 |
| 45 |  | healthy | 140.88 | 150.26 | 119.42 | 79.70 | 51.82 |
| 46 |  | infected | 206.53 | 216.91 | 169.31 | 64.16 | 40.72 |
| 47 |  | infected | 235.74 | 241.38 | 185.05 | 58.70 | 42.32 |
| 48 |  | infected | 220.57 | 227.21 | 171.82 | 58.98 | 43.32 |

### Peroxidase Estimation

Peroxidase (PODs) detoxify the H_2_O_2_ and other hydro peroxides by catalyzing the reduction of ascorbate (AsA), monodehydroascorbate (MDHA), glutathione (GSH) or dehydroascorbate (DHA) depending upon the peroxidase and shielding cells from oxidative damage [27]. PODs were estimated as described by Siddique and co workers in cotton leaf samples as with slight modification. In this assay guaiacol is used which produce brown color upon oxidation [28].

### Reaction Mixture

The reaction mixture was prepared in 50 MM potassium phosphate buffer. The calculated volume of H_2_O_2,_ and guaiacol were added to the buffer to make reaction mixture with final concentrations 40 MM and 20 MM, respectively. The 240 µl of reaction mixture was added to each well of micro well plate and the samples to be estimated were pored to another plate in predefined order. The samples were added to the reaction mixture with the help of multichannel pipette and time was recorded on the mobile stopwatch. Few wells were loaded with only phosphate buffer instead of sample to get the blank values. The absorbance was measured at 470 nm after every 60 sec using ELISA plate reader (BioTek, µ-QuantTM, USA) and 6 reading were recorded.

### Calculations

The PODs activity was estimated by the difference of absorbance. The first minutes readings were used after subtracting from blank reading. The ΔA_470_ /min is the absorbance after first minute minus absorbance of blanks. The following formula was used to calculate enzyme activity in U/mL. Finally the values are expressed as U/g of fresh weight and given in (Table 1).

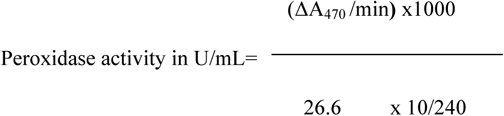

The 26.6 is H_2_O_2_ extinction coefficient, and 10/240 is dilution factor as 10 µl of sample was added to 240 µl of reaction mixture and there are 1000 µl in 1 mL.

### Superoxide Dismutase (SOD) Determination

Superoxide dismutase assay was performed on the basis of NBT reduction inhibition as described by Siddique and coworkers with little modifications [28].

### Reaction mixture and Enzyme Assay

The master reaction mixture was prepared by addition of 50 MM potassium phosphate buffer (pH-7.0), 100 MM Ethylene diamine tetra acetate (EDTA), 1.5 MM Nitro blue tetrazolium (NBT) and 0.1 MM riboflavin in the end according to these ratios 53:4:2:1, respectively. This is light sensitive solution and used immediately after formation. The master mixture was prepared according to the number of samples and each well was loaded with 230 µl master mix. 10 µl of sample was added to each well according to predefined well schedule and few wells were loaded with dH_2_O as control. After addition of all the samples blank reading was recorded using ELISA plate reader (BioTek, µ-QuantTM, USA) at 560 nm wavelength. The ELISA plate was placed in light for the reaction and absorbance was recorder after every 30 sec and took 5 reading. The difference in absorbance was observed to check the pattern of reduction of NBT in controls samples suggests a linear reduction pattern only in first reading after 30 Sec. The enzyme activity was calculated using the following equation to calculate the percentage inhibition by SOD in each sample. The enzyme activity was converted to the weight of the enzyme present in the each gram of sample through appropriate calculation. The values are expressed as U/g of fresh weight and given in (Table 1).

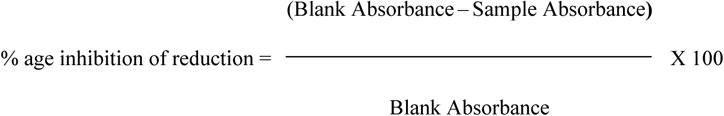

Where, sample absorbance is after 30 sec so the values obtained were multiplied by 2 to obtain 1 minute activity of the enzyme.

### Estimation of Non-Enzymatic Metabolites

The fresh leaves of healthy and virus infected cotton leaves were used to estimate the non-enzymatic antioxidant metabolites (total phenolic compounds and total flavonoid contents).

### Determination of Total Phenolic Contents (TPC)

The TPC of cotton leaves (infected and healthy) collected from different locations were determined as described by [29] with minor modifications. The leave samples (10mg each) were homogenized in 80% acetone in bead beater (Omni Bead Ruptor 24, by Omni International Canada) in 2 mL tubes and homogenates were centrifuged at 10000 rpm for 10 minutes. Supernatant was collected in new eppendorf and use for TPC estimation.

### Reaction Mixture and Standard Curve

The reaction mixture was prepared by dilution 2 mL of Folin-Ciocalteau’s reagent in 16 mL of distilled water. The each well of ELISA plate were loaded with 50 µl of reaction mixture and 5 µl of Gallic acid standers were added and mixed in 3 consecutive wells each to get the mean value for standard curve. The plate was incubated at room temperature for 15 minutes. The 200 µl saturated solution of sodium carbonate (Na_2_CO_3_) was added to the each well and incubated at room temperature for half hour. The absorbance was measured at 750 nm by ELISA plate reader (BioTek, µ-QuantTM, USA).The standard curve was obtain using the mean value of these readings (Supplementary Figure S2). The TPC of cotton leave samples were determined in the same way using 5 µl of supernatant. The value of TPC in our samples linear regression equation (Y= 0.0025 X + 0.0206) obtained from was standard curve was used (Supplementary Figure S2). The values are expressed as mg/g of fresh weight Gallic acid equivalence (GAE) and given in (Table 1).

### Determination of Total Flavonoid Contents (TFC)

Flavonoids are phenolic compounds have antioxidant properties. The flavonoids contents were estimated in the cotton leaves as described by Chanda and Dave with few modifications. The fresh leave samples (10mg each) were homogenized in (200 µl) 50 to 80% ethanol in bead beater (Omni Bead Ruptor 24, by Omni International Canada) in 2 mL tubes and homogenates were centrifuged at 10000 rpm for 10 minutes. Supernatant was collected in new eppendorf and used for TFC estimation and stored in freezer [30].

### Reaction Mixture and Standard Curve

The reaction mixture was prepared by adding 70 µl sample and standard extract each well of ELISA plate followed by addition of 70 µl of sodium Nitrite, NaNO_2_ (5%) and incubated mixture for 5 Minutes. The 15 µl of aluminum chloride, AlCl_3_ (10%) was added to each well and incubated at room temperature for 10 mins followed by addition of addition 70 µl of 1M sodium hydroxide (NaOH) to each well. The plates were incubated for 10 minutes before recoding absorbance at 415 nm by ELISA plate reader (BioTek, µ-QuantTM, USA). Quercetin standers were used to get standard curve (Supplementary Figure S3). The standard curve was obtained using the mean value of these readings. The value of TFC in our samples was determined using linear regression equation (Y= 0.0005X + 0.0055) obtained from standard curve (Supplementary Figure S3). The values are expressed as mg/g and given in (Table 1).

## Results and Discussion

The homeostasis of ROS is dependent on its production and scavenging activity of anti-oxidant enzymes and other anti-oxidant compounds [31]. There are two types of defense mechanism against ROS in nature, enzymatic and non-enzymatic. The enzymatic defense system has variety of scavenger enzymes in plants but there are three main enzymes responsible for enzymatic defense against ROS viz. catalase, peroxidases and superoxide dismutase. The non-enzymatic defense includes compounds with intrinsic antioxidant properties and these compounds are either water soluble such as phenolic compounds, flavonoids, ascorbate glutathione or fat soluble like carotenoids and tocopherols. These antioxidant compounds act as electron donor to reduce ROS to less reactive molecules which are not that harmful to cell. Under stress conditions, the ROS homeostatic defense system becomes disturbed due to increased ROS production. To minimize the damaging effect of ROS, the aerobic organisms up regulate the both enzymatic and non-enzymatic antioxidant defense systems [32]. Present study compared the activity of antioxidant enzymes and some antioxidant compounds under stress caused by cotton leaf curl disease.

### Catalase

Catalase is an antioxidant enzyme present in almost every aerobic cell in abundance and detoxify hydrogen peroxide (H_2_O_2_) a toxic product of aerobic metabolism and ROS produced by pathogens or response to pathogens. This enzyme protect cells from toxicity of H_2_O_2_ which can be lethal if not degraded [33]. It converts two molecules of H_2_O_2_ into water and molecular oxygen [34].

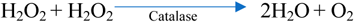

Catalase is tetrameric protein of 240 kDa and each polypeptide of 60 kDa in weight and similar to each other containing a single ferriprotoporphyrin [35]. In the cell it scouts for hydrogen peroxide and have highest turnover number and a single tetramer can convert millions of H_2_O_2_ to water and oxygen in one second. This reaction took place in two steps during first step H_2_O_2_ molecule oxidizes the heme and one porphyrin radical generated and in second step another H_2_O_2_ molecules reduces the porphyrin radical to enzyme resting state producing one molecule of oxygen and two water molecules [36]. In this study the catalase activity in infected cotton leaves is significantly higher than that of healthy leaves as shown in Figure 1 which suggests the increase of ROS in infected cell as compared to the normal and healthy cells.

**Figure 1.**
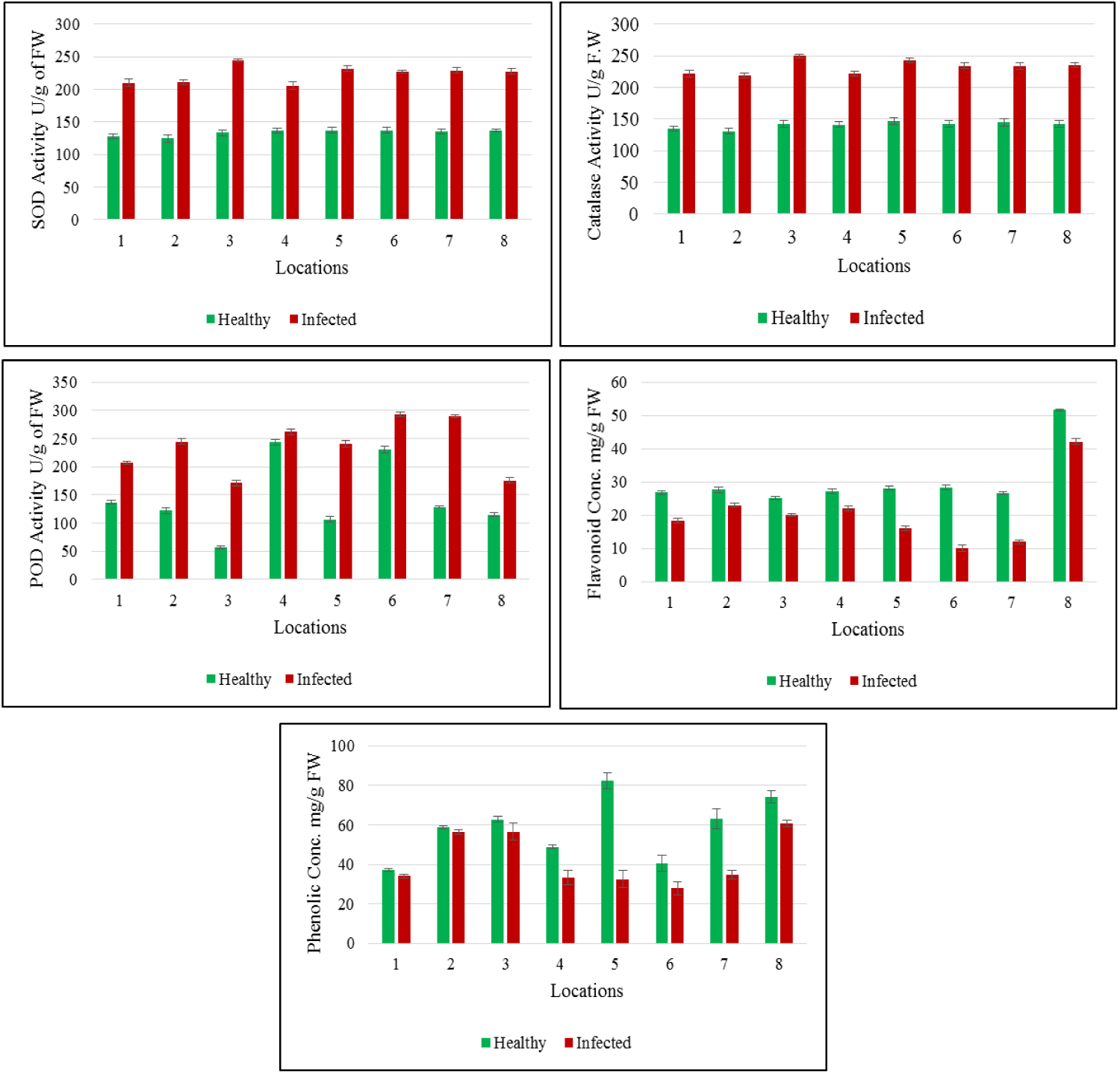
The graphical representation of antioxidant enzymes super oxide dismutase, catalase, peroxidase activity and antioxidant metabolites total flavonoids contents and total phenolic contents in the healthy and CLCuD infected leaves of cotton.

Catalase activity was recorded increased in cotton leaf curl infected leaves and these findings are in agreement with that of Siddique et al. [28]. These finding are in contrast with Riedle-Bauer [37] findings that cucumber mosaic virus does not affect catalase activity in cucumber plants. [38] Ahammed et al. also reported the enhanced activity of catalase under iron stress which is in accordance with our results that the under stress the activity of catalase increases. The increased catalase activity in rice under drought stress is also reported by the [39] which is accordance with our results in this study. [40] Hasanuzzaman et al. also reported the significant increase in catalase activity under abiotic stress which supports our results that the activity of catalase enhanced during stress. These finding suggests that during virus infection the ROS synthesis increases and subsequently plants enzymatic antioxidant defense system also activated to cope with the stress condition.

### Peroxidase

Peroxidases (PODs) are the antioxidant enzymes present in almost every aerobic cell and among the first enzymes respond to phytopathogens in addition to detoxification of hydrogen peroxide (H_2_O_2_) [41]. PODs are involved in Lignification, regulation of cell wall elongation, suberification, wound healing and resistance against phytopathogens [42]. The reaction of PODs are given below

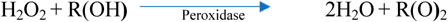

Where, “R” may be Ascorbate (AsA), monodehydroascorbate (MDHA), glutathione (GSH) or dehydroascorbate (DHA) depending upon the peroxidase. Peroxidases counteract the increased hydrogen peroxide along with catalases and plays critical role in the antioxidant defense response of plant [43, 44]. The protective role of peroxidases have been reported against plant disease and tissue injuries [45]. These have greater affinity for H_2_O_2_ than that of catalase and present in cytoplasm and mitochondria. In this study the catalase activity in infected cotton leaves is significantly higher than that of healthy leaves as shown Figure 1 which suggests that, this group of enzymes have role in plant defense against pathogens.

These finding of enhanced PODs activity in infected cells is in accordance with the finding of [46] who observed immediate increase of PODs level after inoculation of virus. [47] also observed increased PODs activity after inoculating tomato leaves with *A. alternate* leaf blight causing pathogen which is also in accordance with our results the higher expression of PODs after infection. Our results are also in agreement with [28] for the increased activity of PODs in plants grafted with virus infected scion. The increased PODs activity in rice under drought stress is also reported by the [39] which is accordance with our results in this study. These finding suggests that during virus infection the activity of PODs also increases as defense against pathogens.

### Superoxide Dismutase (SOD)

Superoxide dismutase are the metalloproteins which catalyze the dismutation of superoxide (O_2_^-^) radicals to molecular oxygen or H_2_O_2_. The SODs are classified into three types according to the metal ion association and important part of cells antioxidant defense system [48]. The H_2_O_2 -_ produce during this reaction is catalyzed to water and oxygen by catalases and peroxidases. The reaction scheme is given below.

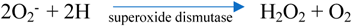

SODs are the first line defense of cell against ROS and also plays important role against the pathogenic infections. The SOD activity in infected cotton leaves is significantly higher than that of healthy leaves as shown in Figure 1. These findings suggests that, this enzyme have role in plant defense against pathogens. These finding of enhanced SOD activity in infected cells is in contrast with the finding of [46] who observed decrease of SODs activity after inoculation of virus and with [28] for the increase activity of SODs in plants grafted with virus infected scion. These findings are in accordance with [39] who reported increased SOD activity in rice under drought stress. [40] also reported the significant increase in SOD activity under abiotic stress which supports our results that the activity of SOD enhanced during stress. We can conclude that SOD activity altered during stress.

### Total Phenolic Contents (TPC)

Polyphenols of plants are distinct group of phenolic compounds (phenolic acids, flavonoids, stilbenes, lignins etc.) having ideal structure for scavenging of free radicals. Their property of electron or hydrogen donation makes them good antioxidant compounds [49]. These phenolic compounds enhance the mechanical strength of cell wall of the host cells may be by synthesis of lignin and suberin and form a physical barrier for the pathogens [50, 51]. In this study the amount of Phenolic compounds is significantly reduced in infected plants shown in Figure 1. A significant difference of total phenolic contents observed between the samples from different locations this might be due the difference of cultivars or climatic conditions affected accumulation of these compounds. These results are in accordance with the results of [28] who reported that reduction of phenolic compounds in infected/inoculated cotton cultivars as compared to the healthy/non inoculated plants. Chatarjee [52] reported that virus infected Mesta plants have lower total phenolic contents as compared to that of healthy plants and also observed that the resistant cultivars have high level of total phenolic compounds as compared to the susceptible varieties the first part is in accordance with our results of this study. (Chaubey & Mishra, 2020) also reported that the leaf curl disease to chili also reduced the total phenolic contents in the infected leaves as compared to the healthy leaves which is in agreement with our findings in this study. It can be concluded that in the infected leaves the level of total phenolic compounds drop significantly as compared to that of healthy plants. Moreover, the climatic condition also effect the level of these compounds.

### Total Flavonoid Contents (TFC)

Flavonoids are phenolic compounds have antioxidant properties and can scavenge free radicals as well as chelate the metal ions (Iron, copper etc.) subsequently inhibiting the enzymes responsible for generation of free radicals [53]. These compounds occur in the form of glycosides in plants and hypothesized to perform several roles like photoproduction, hormonal translocation modulation and sequestration of ROS [54]. Flavonoids can scavenge all known ROS depending on their structure. During stress condition the level of TFC may be altered as in our study. The level of flavonoids in CLCuD infected plants has dropped little as compared to healthy leaves as shown in Figure 1. There might be difference of verities which express different flavonoid contents and the information of the cotton varieties is not available. The difference might be due to the different climatic condition as the samples were collected from different locations. The overall results shows that the flavonoid content in the infected leaves is lower than that of healthy leaves from the same field. These results are in accordance with that of [28] who perform the assay for the cotton leaf curl disease infected plants and healthy plants in addition to that of disease resistant cotton cultivars which express more flavonoid contents than that of susceptible cultivars. It can be concluded that in the infected leaves the level of total flavonoid contents drop significantly as compared to that of healthy plants. Moreover, the climatic condition also effect the level of these compounds.

### Conclusion

These findings suggests that antioxidant enzymes catalase, peroxidases and superoxide dismutase along with antioxidant compounds constitue the first line of defense with key role in total defense machanism of biological system. The activity of all antioxidant enymes during cotton leaf curl disease increases and the level of antioxidant compounds decreased.

## Supplementary Data

**Figure S1.**
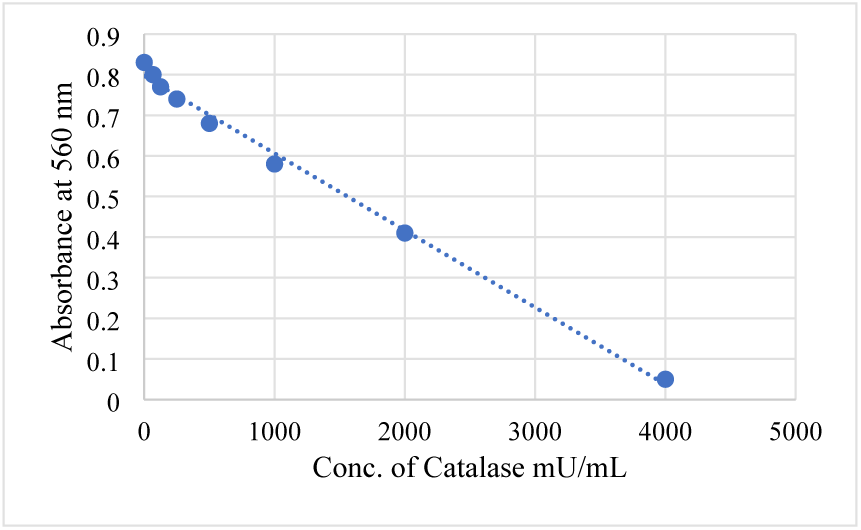
The Standard Curve for Catalase activity

**Figure S2.**
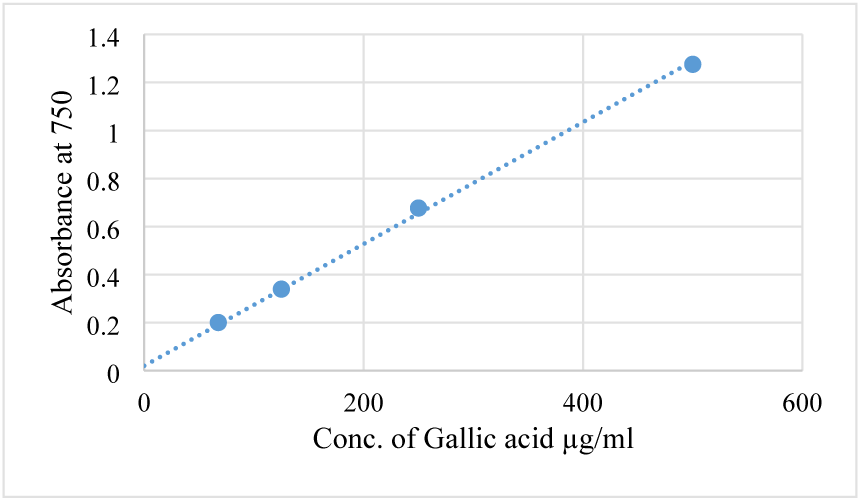
The Standard Curve for Gallic acid (GA) for quantification of total phenolic contents.

**Figure S3.**
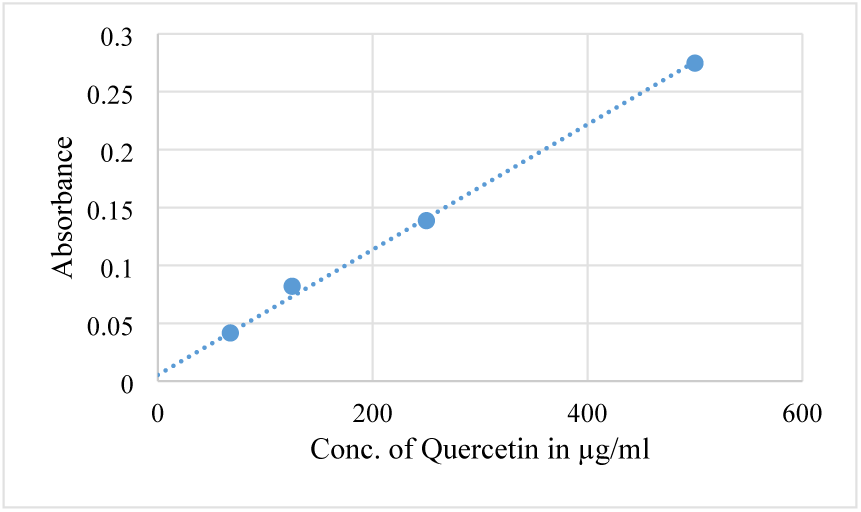
The Standard Curve for Quercetin for quantification of total flavonoid contents

